# On the relationship between the rate of learning and retrieval-induced forgetting

**DOI:** 10.1101/350959

**Authors:** Ádám Markója, Ágnes Szőllősi, Mihály Racsmány

## Abstract

A plethora of studies demonstrated that repeated selective retrieval of target items from semantic categories has an adverse memory effect on semantically-related memories, a phenomenon called retrieval-induced forgetting (*RIF*). However, there is a range of boundary conditions for RIF. For instance, forming interconnections between target and competitors, long-term delay without sleep between practice and final recall, and the form of learning all attenuate the effect of selective practice on the accessibility of semantically-related competitors. The aim of the present research was to investigate the latent general preconditions behind the reductions of *RIF*. In Experiment 1 participants learned category-exemplar pairs with repeated study or combined study-full test sessions followed by a selective retrieval practice and a full cued-recall test. We found lower difference between performance on the non-practiced items from practiced categories (Rp-) and their baseline from unpracticed categories (Nrp-) suggesting a reduction in retrieval-induced forgetting. However, regression analysis revealed that this was possibly caused by the increased recall performance independently from the presence of an initial full retrieval session. Therefore, in Experiment 2 participants learned the same pairs through a study followed by two study, two test or combined study-test cycles. As the consequence of increased rate of learning we confirmed the complete absence of retrieval-induced forgetting with Bayes factor analysis. Our results suggest that the adverse memory effect of selective retrieval practice shows a non monotonic dependency on the strength of the mnemonic representations.

## Introduction

Although there are many advantages of retrieval-based learning, even in real life (Roediger III & Karpicke, 2006), repeated retrieval attempts also have an adverse effect on semantically related, non-practiced items, which were studied before selective retrieval practice (Anderson, Bjork, & Bjork, 1994). This effect, called retrieval-induced forgetting (RIF), was first demonstrated by the so-called retrieval practice paradigm. In the original task (Anderson et al., 1994) participants learned a set of category-exemplar pairs (e.g., Fruit – Orange). Then, in a selective retrieval practice phase, participants repeatedly recalled half of the members from half of the categories during cued recall trials (e.g., Fruit – Or__). As a detrimental consequence of selective target strengthening, the authors observed reduced recall performance for the non-practiced items from the practiced categories, compared to recall performance for items from the non-practiced categories.

Retrieval-induced forgetting, as a general phenomenon in the literature of forgetting, has several key features and boundary conditions. One of these key features is that RIF is retrieval-specific, i.e., RIF does not occur when participants has an extra selective study opportunity (Ciranni & Shimamura, 1999; Experiment 5) or applied a reversed retrieval practice^1^ (Anderson, Bjork, & Bjork, 2000). Additionally, in the original experiment, Anderson et al (1994) manipulated the target and competitor strength and found that target strength had no impact on RIF, while there was no RIF effect when competitors were week. Moreover, there has been no correlation between the target strengthening and RIF, indicating the strength independent nature of RIF (Hulbert, Shivde, & Anderson, 2012). Based on these features, along with the interference dependent and cue independent nature of RIF, the inhibition based account became the most dominant account in the RIF literature, which assumes an active suppression mechanism beside interference and associative blocking, implicating the involvement of the prefrontal cortex (Anderson, 2005; Anderson & Levy, 2007).

While the inhibition based account specified important characteristics of RIF, contradictory results can be found which cannot be completely explained by an inhibition account (e.g. Jakab & Raaijmakers, 2009; Jonker & MacLeod, 2012; Jonker, Seli, & MacLeod, 2012; Raaijmakers & Jakab, 2013). Earlier, Norman, Newman, & Detre (2007) introduced a neural network model of RIF based on oscillated inhibition mechanisms. Along with the semantic memory (cortical) layers their model also involved a hippocampal layer which was able to follow the contextual changes in accordance with the demand on implementation episodic-based retrieval. Their simulations suggested that RIF is a context-dependent phenomenon, leading to a whole new family of explanations based on the crucial role of context shift between the study and selective retrieval practice (Jonker, Seli, & MacLeod, 2013, 2015). Additionally, the predictions of their neural network model challenged the universality of the strength-independent nature of RIF suggesting a non-monotonic relationship between the magnitude of retrieval-induced forgetting and target strength (Norman et al., 2007; see Simulation 2.2). These predictions for the high target strength had an experimental confirmation through a peculiar RIF task containing arithmetic operations (Campbell & Phenix, 2009; for a non-monotonic relationship between strength and RIF see also Keresztes & Racsmány, 2013).

Results supporting the context-based account of RIF pose another challenge to a solely inhibition-based accounts. Recent studies have demonstrated that involving one single initial full test before the selective retrieval practice phase leads to a context update of the study phase and leads to the lack of retrieval-induced forgetting (Racsmány & Keresztes, 2015).

From the point of view of strength-independent nature of RIF contradictory results demonstrated that retrieval-induced forgetting can be reduced either by manipulating the target strength (Campbell & Phenix, 2009; Norman et al., 2007) or the context of encoding (Jonker & MacLeod, 2012; Jonker et al., 2012; Racsmány & Keresztes, 2015). Additionally, it has been already demonstrated that involving a 12-hour retention interval between selective practice and final recall (with no sleep during the delay) moderates the effect or retrieval-induced forgetting (Abel & Bäuml, 2012; Racsmány, Conway, & Demeter, 2010; but see MacLeod & Macrae, 2001), as well as using an integrated set of representations (Anderson & McCulloch, 1999). Altogether, these findings suggest that retrieval-induced forgetting can be interpreted as existing on a continuum influenced by multiple components than as an all-or-nothing effect.

Many of these factors (especially the delay and the strategy during encoding) may be contributing not only to the adverse and favourable effects of selective retrieval practice rather to the entire encoding and retrieval procedure. This framework suggests that retrieval-induced forgetting may be determined by a more general factor shaped through many experimental manipulations, namely the rate of learning before selective retrieval practice. As a consequence, our goal was to examine the multidirectional relationship between the rate of learning determined by overall recall success, target strength, and retrieval-induced forgetting. We designed an experimental procedure based on the Experiment 2 and 3 of Racsmány & Keresztes (2015). As a modification, we included a longer delay between the initial test and the following selective retrieval practice. Moreover, neither additional delays nor reexposure blocks were applied between the individual practice cycles. These modifications were necessary to make the results directly comparable to the conventional RIF experiments (e.g. Anderson et al., 1994) and reduce the confounding factors during the competition process observed in the selective retrieval practice. Additionally, to counterbalance these changes in global representational strength we administered a control condition involving an extra study after the initial study phase. Based on the supportive results from this manipulation we conducted another experiment to investigate the high strength-related limitations suggested by Norman et al. (2007) in the context of not only the target strengthening, rather the global rate of learning.

## Experiment 1

### Materials and Methods

#### Participants

Eighty participants (14 men; age range: 19-28 years, *M* = 20.7, *SD* = 1.9) were recruited at Eötvös Loránd University, Budapest, Hungary. Participants were randomly assigned to an extra study (7 men; age range: 19-26 years, *M* = 20.7, *SD* = 1.7) or a full test group (7 men; age range: 19-28 years, *M* = 20.7, *SD* = 2.1). Subjects received extra course credits for their participation.

Both experiments was approved by the Ethical Committee of the Budapest University of Technology and Economics, Hungary. All participants gave written informed consent.

#### Design and Materials

Stimuli were 48 words selected from eight semantic categories (six items per category). Out of these 48 words, 12 words from two categories were fillers (six items per category). For each subject, a pseudo-randomized selection method was used to assign the remaining 36 items from six categories into one of the four item types. Three categories were practiced (Rp). Half of the words belonging to these categories were practiced items (Rp+, three items per category), while the other half of the words were the unpracticed items from the practiced categories (Rp-, three items per category). The remaining words were the non-practiced baseline items (Nrp). Baseline items were randomly assigned into either of two conditions. Half of the words were baseline items for the practiced words (Nrp+), whereas the remaining words were baseline items for the unpracticed words (Nrp-).

#### Procedure

Our experiment consisted of four main phases: an initial study phase, an extra study or a full test phase (varied between participants), a selective retrieval practice phase consisting of 3 cycles, and a final test phase. Immediately before the initial study phase, participants were instructed to memorize the words (i.e. the category exemplars) together with their category label (also presented on the computer screen), in order to recall the exemplars in response to the category labels at the following memory test. During the initial study phase, participants were presented all 48 category-word pairs. The category label was presented on the left side of the computer screen, whereas the category exemplar was presented on the right side of the computer screen. Participants saw only one category-exemplar pair at a time and each pair remained on the screen for 5000 ms, followed by a 500-ms inter-stimulus interval (ISI). The stimuli were presented in a pseudo-random order with two restrictions: presentation started and ended with two fillers, and three subsequent trials never included items from the same category.

The initial study phase was followed by either another study phase (extra study group) or an initial retrieval phase (full test group). In the extra study group, all category-exemplar pairs were presented again. Stimulus presentation time and ISIs were the same as in the initial study phase. In the full test group, participants were asked to recall the 48 exemplars in response to the category labels. In each trial of the initial retrieval phase, a category label was shown on the left side of the screen, and the first two letters of the target item on the right side of the screen. Participants were instructed to recall the target word and to fill in the missing letters by using a standard Hungarian keyboard. The category label and the first two letters of the target word were on the screen either until a response was entered or for 8000 ms (with an ISI of 500 ms). This phase was followed by a series of arithmetic distractor tasks in both groups. The arithmetic tasks were additions and extractions with three- or four-digit numbers, and each arithmetic task was followed by a feedback in the form of the correct answer was presented with the participant.

Immediately after the arithmetic distractor tasks, a selective retrieval practice phase followed. The selective retrieval practice phase consisted of three cycles. Participants’ task in both groups was the same as the full test group’s task was in the initial retrieval phase. However, only half of the words from half of the categories (nine Rp+ items with six fillers) were practiced. Each practice cycle started and ended with two filler items. Then, participants performed a series of arithmetic distractor tasks for eight minutes.

In the final cued recall test phase, we tested subjects’ memory for all stimuli. Participants’ task was the same as the full test group’s task was in the initial retrieval phase. However, we used two modifications: participants were given only the first letter of the target word together with the category label, and this test phase consisted of two blocks in order to control output interference (see Anderson, 2003). In the first block, participants were required to recall the Rp- items pseudo-randomly intermixed with their baseline (Nrp-) items, and in the second block, they were required to recall the Rp+ items pseudo-randomly intermixed with their baseline (Nrp+) items. There was no delay between these two blocks, and both blocks started and ended with two filler items. The experimental procedure is illustrated in Figure 1.

**Figure 1.**
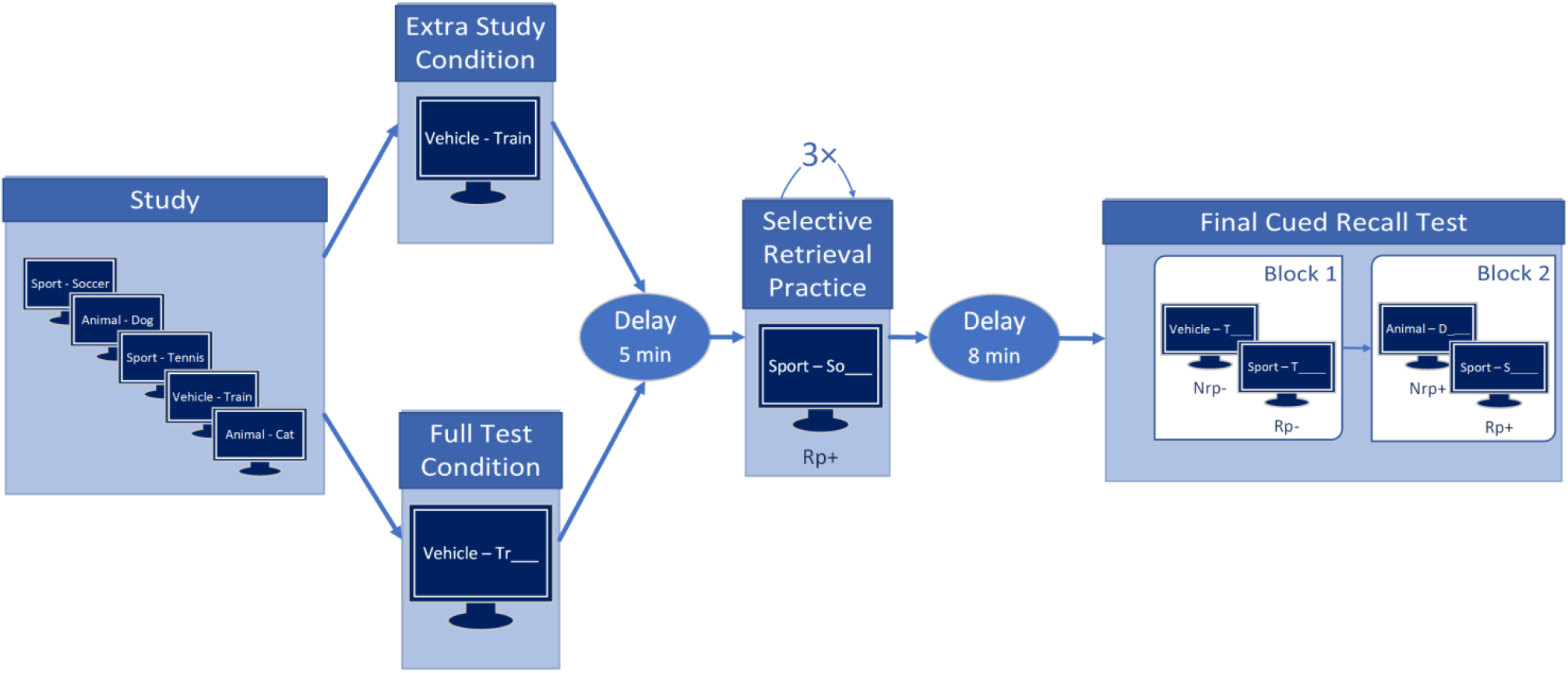
Schematic experimental procedure for Experiment 1.

Stimulus presentation and randomization were performed with PsychoPy (version 1.84) and NumPy (version 1.12) Python (version 2.7) modules.

### Results and discussion

The dependent variable was recall performance (percentage of correctly recalled items per condition). The data was screened for outliers (outliers were defined as the percentage of correctly recalled items is more than three standard deviations away from the group mean in each condition) and none of the participants were identified as outlier.

To reliably confirm the presence or the absence of effects we computed Bayes factors together with Null Hypothesis Significance Testing in both experiments. For linear models (regression, BANOVA) we used uninformed prior distribution for H_1_ assuming standardized effect size comes from *Cauchy* distribution with scale parameter of √2/2. To test whether RIF occurs within conditions we also used *Cauchy*(*r* = √2/2), while practice effect expected from a wider range: *Cauchy*(*r* = √2) suggested by our previous studies (see supplementary material).

#### Recall performance

We conducted a 4×2 mixed effect ANOVA with item TYPE (Rp+, Rp-, Nrp+, and Nrp-) as a within-subject variable and CONDITION (full test vs. extra study) as a between-subject variable. Item TYPE had a significant main effect on recall performance, F(3,234) = 28.47, *p* < .001, *MSE* = 177.09, η^2^_p_ = .27, while there was no main effect of CONDITION. However, the interaction between TYPE and CONDITION was significant, *F*(3,234) = 2.99, *p* = .03, *MSE* = 177.09, η^2^_p_ = .03. According to the Bayesian ANOVA (BANOVA) the best fitting model was the one containing item TYPE and TYPE ×CONDITION interaction term with a Bayes factor of 6.51 × 10^12^ relative to null model (Bayes factor for the other models can be seen in Table 1). We compared all other models to the highest fitting model. The posterior probability of the interaction model has a Bayes factor of 1.19, again the model containing only item TYPE, therefore the simple main effect model and interaction model has about equal posterior probability. Besides, interaction model has more than five times higher posterior probability in comparison to others therefore all the rest alternative models can be excluded (see Table 2).

**Table 1.** Bayes factors for each possible BANOVA model for Experiment 1 AGAINST NULL model.

| Model | Bayes factor |
| --- | --- |
| **Type + Type×Condition + Participant** | **$6.51 \times 10^{12}$** |
| **Type + Participant** | **$5.49 \times 10^{12}$** |
| Type + Condition + Type×Condition + Participant | $1.22 \times 10^{12}$ |
| Type + Condition + Participant | $1.10 \times 10^{12}$ |
| Type×Condition + Participant | 0.47 |
| Condition + Participant | 0.17 |
| Condition + Type×Condition + Participant | 0.08 |

**Table 2.** Bayes factors for each possible BANOVA model for Experiment 1 IN FAVOUR OF THE STRONGEST model.

| Model | Bayes factor |
| --- | --- |
| **Type + Type×Condition + Participant** | **1.00** |
| **Type + Participant** | **1.19** |
| Type + Condition + Type×Condition + Participant | 5.34 |
| Type + Condition + Participant | 5.92 |
| Type×Condition + Participant | $1.37 \times 10^{13}$ |
| Condition + Participant | $3.83 \times 10^{13}$ |
| Condition + Type×Condition + Participant | $8.02 \times 10^{13}$ |

During the upcoming contrast analyses in both experiments we will use contrast weights to separate the the effect of practice (difference between recall performance on the Rp+ and Nrp+ items) from the effect of retrieval-induced forgetting (RIF, difference between recall performance on the Rp- items and Nrp- items) in order to conduct general linear hypothesis tests (GLHT) on estimated marginal means, using Holm (1979) correction method and we also perform pairwise Bayesian t-tests.

We found significant difference between the Rp+ items: M = 81.8, SD = 14.1 and Nrp+: M = 67.1, SD = 16.0.96 items, therefore we can confirm the practice effect independently from conditions: t(234) = 7.00, *p* < .001, *β* = 14.7 [9.99, 14.46], with a Bayes factor of 3.8×10^6^, and also found significant difference between Rp-: M = 63.5, SD = 17.9 and Nrp- items.: M = 71.5, SD = 14.7 items, which implicates the presence of retrieval-induced forgetting: *t*(234) = −3.83, *p* < .001, *β* = −8.06 [−12.79, −3.32]. with a Bayes factor of 112.2.

We conducted simple effect analysis to investigate the effect of item type within each group separately. We observed significant practice *t*(234) = 6.81, *p* < .001, *β* = 20.28 [13.58, 26.97]; *BF* = 6.0×10^4^ and RIF, *t*(234) = −3.45, *p* < .001, *β* = −10.28 [−16.97, −3.58]; *BF* = 62.7 effects within the extra study group; within the full test group, however, only the practice effect was significant, *t*(234) = 3.08, *p*=.004, *β* = 9.17 [2.47, 15.86] with a Bayes factor of 15.4, while the RIF effect was marginally significant, *t*(234) = −1.96, *p* = .051, *β* = −5.83 [−12.53, 0.86] with a Bayes factor of 0.84, which is a very weak support for the absence of retrieval-induced forgetting (see Figure 2). Additionally, the decrement in Practice effect can be seen on the contrast between Bayes factors is more likely caused by the baseline difference (recall on Nrp + items compared between extra study condition and full test condition): *t*(253) = −2.141, *p* = .033, *β* = −7.5 [−14.40, −0.6]; *BF* = 1.68 than the difference on practiced (Rp +) items between groups: *t*(253) = −0.396, *p* = .692, *β* = 3.6 [−3.2, 10.51], *BF* = 0.41.

**Figure 2.**
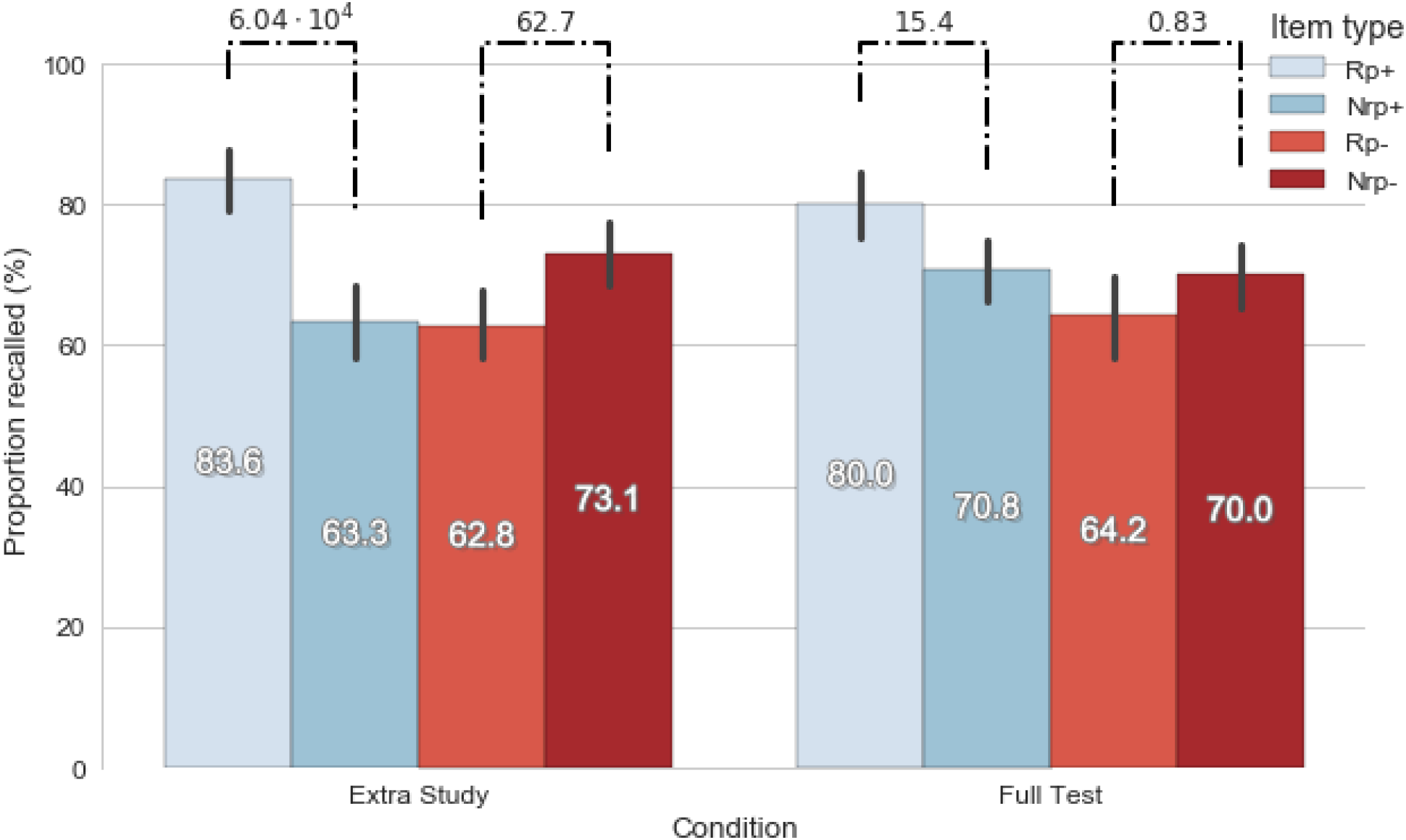
Performance on each item TYPE compared between groups during Experiment 1. Bayes factors can be seen for the differences between recall rate on item TYPE pairs per CONDITION against the NULL. Error bars represent 95% confidence intervals.

#### Regression models of retrieval-induced forgetting

We compared four regression models to determine whether condition, overall performance and/or practice predict the emergence of retrieval-induced forgetting. We calculated RIF scores (difference between recall performance on Nrp- and Rp- items), practice scores (difference between recall performance on Rp+ and Nrp+ items) and overall recall performance scores (mean of correct responses regardless of item type).

In the first model, RIF scores were regressed on practice scores (*model1*: RIF ~ Practice). The fit of this model was not significant, *F*(1, 78) = 1.03, *p* = .314.

Next, overall recall performance was added and the new model (*model2*: RIF ~ Practice + Recall performance) had a significant fit, *F*(2, 77) = 4.22, *p* = .018 with an R^2^ of .099 (R^2^_adj_ = .075). In this model, recall performance negatively predicted the RIF scores (*β* = −0.61, *p* = .008, see Figure 3), while practice score did not predict the RIF scores (*β* = 0.096, *p* = .348). Model fit significantly improved in comparison to *model1, F*(1, 77) = 7.66, *p* = .007, R^2^_change_ = .086).

**Figure 3.**
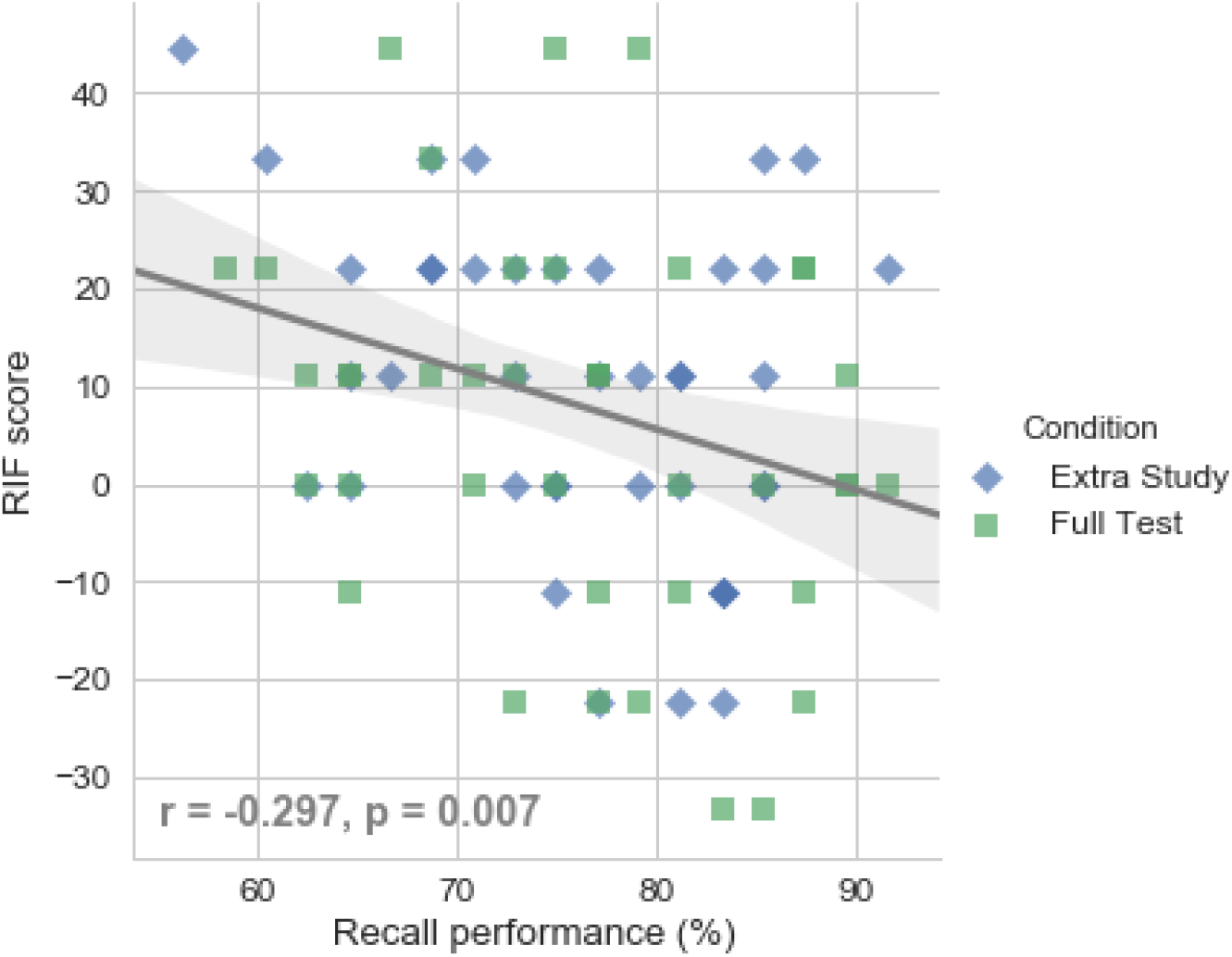
The connection between overall recall performance and the occurrence of retrieval-induced forgetting during Experiment 1. *r* and p values refer to the result of Pearson correlation test.

In the next step, the interaction term was entered to the model (*model3*: RIF ~ Practice + Recall perf. + Practice × Recall perf.). Relative to model2, the fit significantly improved, *F*(1, 76) = 4.54, *p* = .036] and the model fit was significant, *F*(3, 76) = 4.46, *p* = .006 with an R^2^ of .150 (R^2^_adj_ =.116). Recall performance was not a significant predictor in the *model3*, while practice score (*β* = 1.l92, *p* =.029) and the practice×recall interaction (*β* = −2.479, *p* = .036) had a significant effect on RIF scores.

In order to analyse and illustrate the interaction, we divided participants into low, moderate and high performance subgroups based on the quartiles of overall recall performance (see Figure 4). The regression curve only had positive slope in low recall range while we did not observe any connection between practice and RIF scores in moderate and high ranges, see Figure 5.

**Figure 4.**
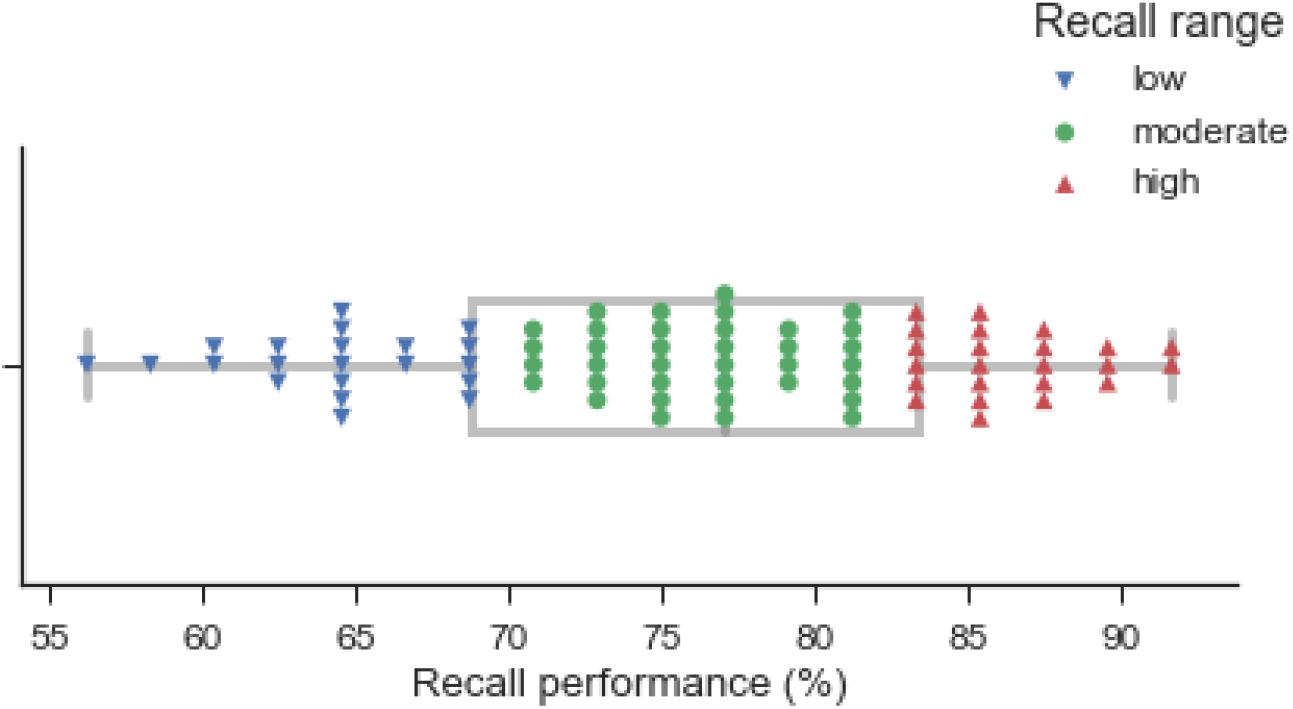
Quartile-based recall ranges in Experiment 1.

**Figure 5.**
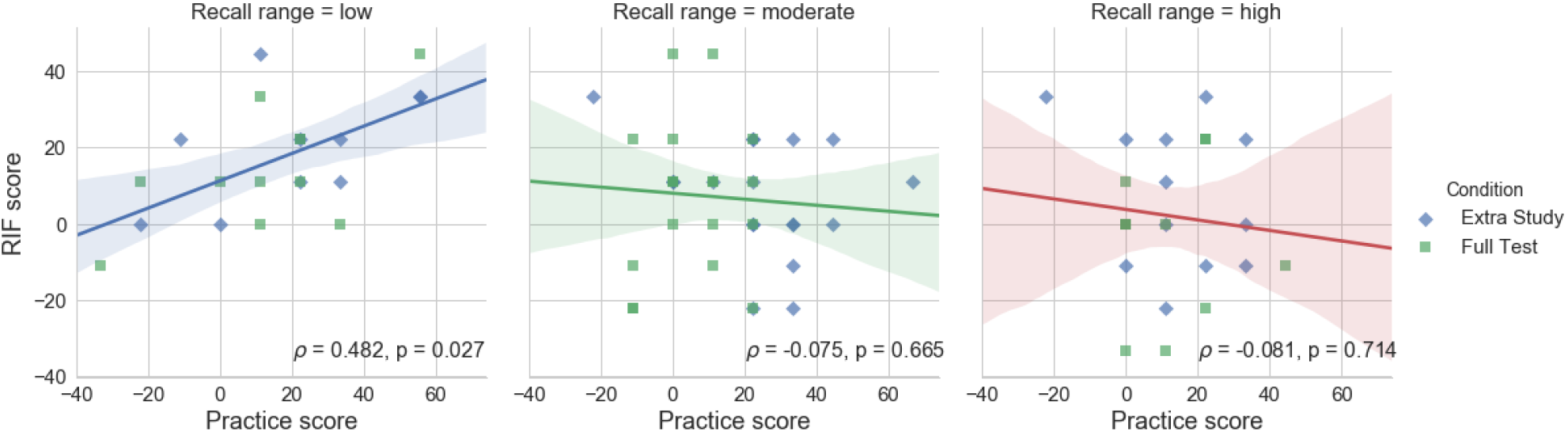
Connection between RIF scores and practice effect within recall ranges in Experiment 1. *ρ* and p values on figure refer to the results of Spearman rank correlation test on each subset.

In the fourth model, we entered condition as a categorical predictor (*model4*: RIF ~ Practice + Recall perf. + Condition + Practice×Condition + Recall perf.×Condition + Practice×Recall perf. + Practice×Recall. Perf.×Condition), but the new model did not fit better than *model3, F*(4, 72) = 0.97, *p* = .43.

Bayesian linear regression analysis also suggests that only *model2* (without the practice score as predictor): *BF* = 5.72 and *model3*: *BF* = 5.53 are evidently capable to explain the variance of RIF scores better than the null model (see Table 3) with approximately equal posterior probabilities suggested by Bayes factor of 1.03 in favour of *model2* over *model3* (see Table 4).

**Table 3.** Bayes factors for each possible linear regression model on RIF scores AGAINST THE NULL model.

| Model | Bayes factor |
| --- | --- |
| **Recall_perf.** | **5.72** |
| **Recall_perf. + Practice + Recall_perf.×Practice** | **5.53** |
| Recall_perf. + Practice | 2.42 |
| Recall_perf. + Practice + Recall_perf.×Practice + Condition | 2.28 |
| Recall_perf. + Condition | 2.27 |
| Recall_perf. + Practice + Recall_perf.×Practice + Condition<br>+ Practice×Condition | 1.84 |
| Recall_perf. + Practice + Recall_perf.×Practice + Condition<br>+ Recall_perf.×Condition | 0.84 |
| Recall_perf. + Practice + Condition | 0.83 |
| Recall_perf. + Practice + Condition + Practice×Condition | 0.77 |
| Recall_perf. + Practice + Recall_perf.×Practice + Condition<br>+ Recall_perf.×Condition + Practice×Condition | 0.74 |
| Recall_perf. + Condition + Recall_perf.×Condition | 0.67 |
| Recall_perf. + Practice + Recall_perf.×Practice + Condition<br>+ Recall_perf.×Condition + Practice×Condition +<br>Recall_perf.×Practice×Condition | 0.45 |
| Condition | 0.39 |
| Practice | 0.36 |
| Recall_perf. + Practice + Condition +<br>Recall_perf.×Condition + Practice×Condition | 0.30 |
| Recall_perf. + Practice + Condition +<br>Recall_perf.×Condition | 0.29 |
| Practice + Condition | 0.12 |
| Practice + Condition + Practice×Condition | 0.10 |

**Table 4.** Bayes factors for each possible linear regression model on RIF scores IN FAVOUR OF THE STRONGEST model.

| Model | Bayes factor |
| --- | --- |
| **Recall_perf.** | **1.00** |
| **Recall_perf. + Practice + Recall_perf.×Practice** | **1.04** |
| Recall_perf. + Practice | 2.37 |
| Recall_perf. + Practice + Recall_perf.×Practice + Condition | 2.51 |
| Recall_perf. + Condition | 2.52 |
| Recall_perf. + Practice + Recall_perf.×Practice + Condition<br>+ Practice×Condition | 3.11 |
| Recall_perf. + Practice + Recall_perf.×Practice + Condition<br>+ Recall_perf.×Condition | 6.82 |
| Recall_perf. + Practice + Condition | 6.93 |
| Recall_perf. + Practice + Condition + Practice×Condition | 7.43 |
| Recall_perf. + Practice + Recall_perf.×Practice + Condition<br>+ Recall_perf.×Condition + Practice×Condition | 7.69 |
| Recall_perf. + Condition + Recall_perf.×Condition | 8.58 |
| Recall_perf. + Practice + Recall_perf.×Practice + Condition<br>+ Recall_perf.×Condition + Practice×Condition +<br>Recall_perf.×Practice×Condition | 12.68 |
| Condition | 14.81 |
| Practice | 15.77 |
| Recall_perf. + Practice + Condition +<br>Recall_perf.×Condition + Practice×Condition | 18.96 |
| Recall_perf. + Practice + Condition +<br>Recall_perf.×Condition | 19.54 |
| Practice + Condition | 49.73 |
| Practice + Condition + Practice×Condition | 59.50 |

Using a between subject manipulation of learning type we found reduction in the amount of retrieval-induced forgetting when participants memory was strengthened with a full cued recall test before the selective retrieval practice in comparison with participants who learned category-item types through two study phases suggested by the significant interaction in ANOVA model and the results on subgroups. Regression analysis revealed that the observed reduction can be rather caused by the increased arte of learning and the moderated connection between selective retrieval practice and RIF.

## Experiment 2

### Materials and Methods

#### Participants

Ninety participants (21 men; age range: 18-28 years, M=20.83, SD=2.01) were recruited at Eötvös Loránd University. Participants were randomly assigned to a double extra study („S-S-S”, 7 men; age range: 18-28 years, *M* = 20.8, *SD* = 2.1), an extra study - full test („S-S-T”, 7 men, age range: 18-26 years, M = 21.2, *SD* = 2.1) or a double full test („S-T-T”, 7 men, age range: 18-24 years, *M* = 20.5, *SD* = 1.8) condition.

#### Design, Materials and Procedure

Stimuli (category-exemplar pairs) and its assignment into one of the four previously described item types (Rp+/-, Nrp+/-) were identical to Experiment 1. However, we applied two modifications on the procedure. First, between the study and the selective retrieval practice phases we used two additional studies or test phases instead of one resulting three different conditions varied between subjects. In the first group (called „S-S-S” as **S**tudy-**S**tudy-**S**tudy) participants had two extra study phases (they were presented with all of the category-exemplar pairs same as Experiment 1 a total of three times) before selective retrieval practice. In the „S-S-T” (**S**tudy-**S**tudy-**T**est) condition participants had one extra study and one full initial test (identical to Experiment 1: they had to recall all the previously studied items with the category cue and the first two letters). In the third group, called „S-T-T” (**S**tudy-**T**est-**T**est) participants had two full tests after the study phase. The next modification was, the last study or test phase was immediately followed by the selective retrieval practice phase (as described in the Experiment 1) without any delay. In each group, this practice phase was followed by an 8-minute delay while participants were given arithmetic distractor tasks and after that, participants had a final cued recall test, same as in Experiment 1.

As we can see above, we varied only the order and the number of phases, while stimuli and runoff (item counts, presentation times, inter-stimulus interval) of the particular phases (study, initial full test, selective retrieval practice and final cued recall test) remained identical to Experiment 1. The experimental procedure is illustrated in Figure 6.

**Figure 6.**
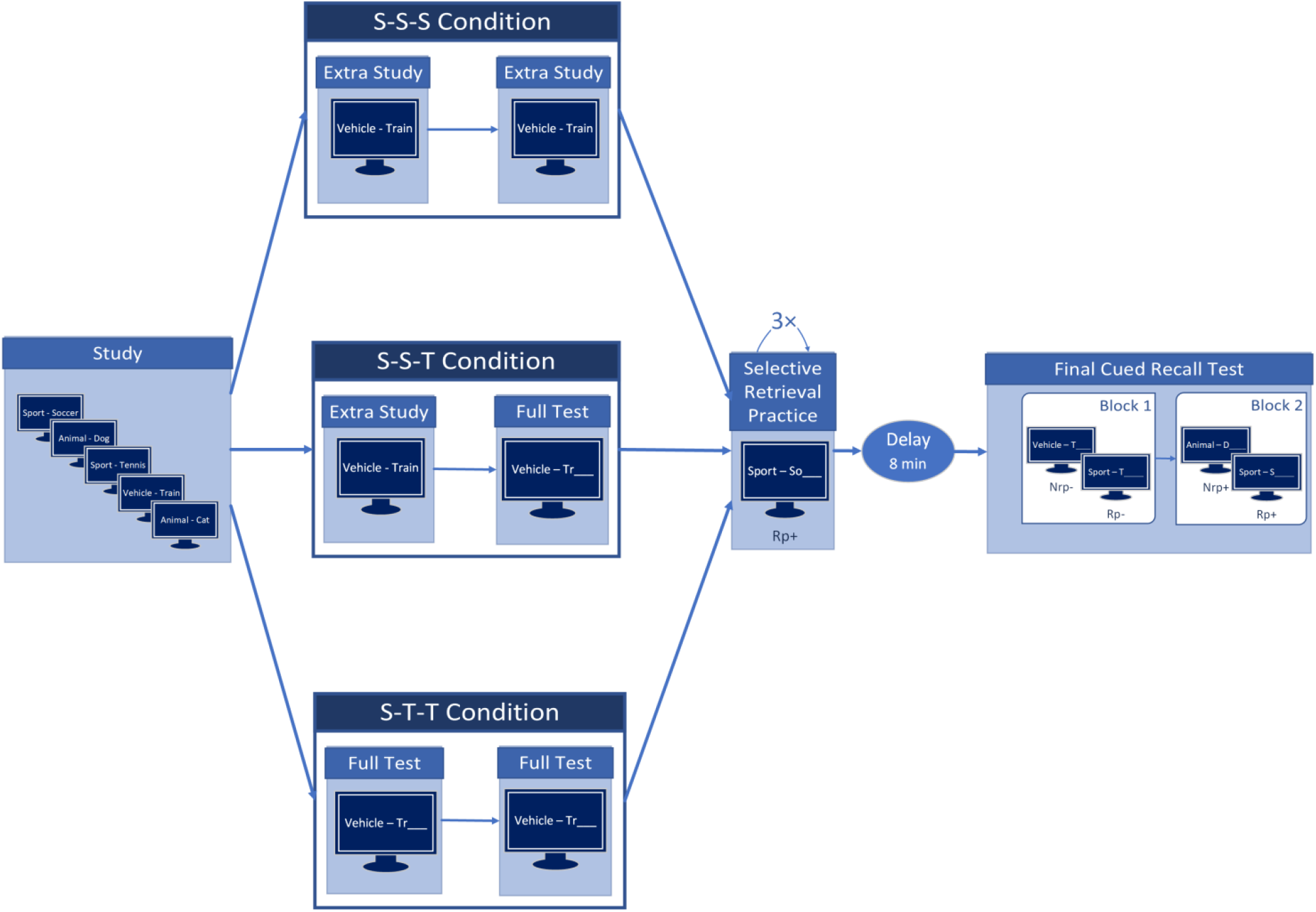
Schematic experimental procedure for Experiment 2.

**Figure 7.**
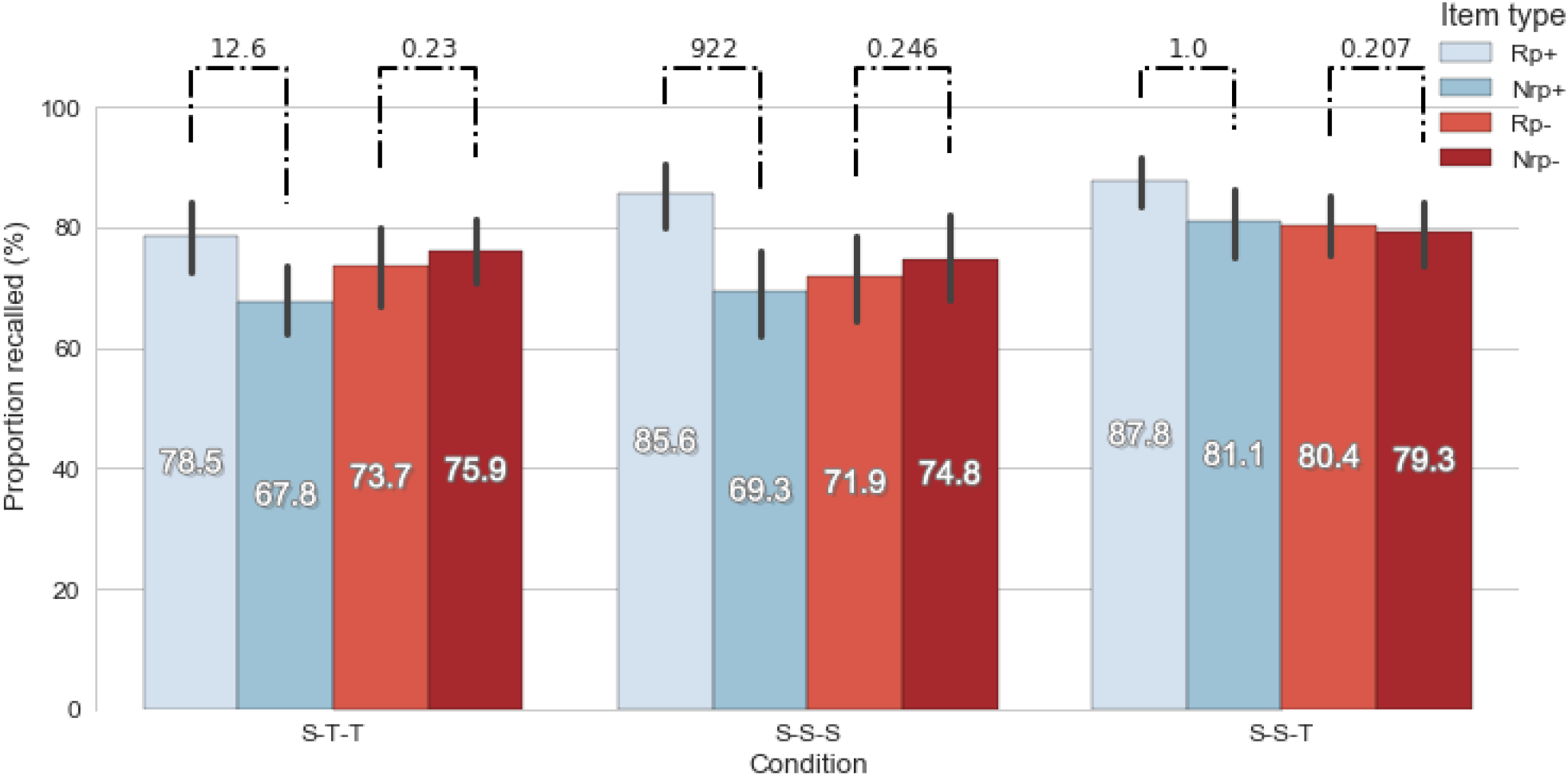
Performance on each item TYPE compared between groups during Experiment 2. Bayes factors can be seen for the differences between recall rate on item TYPE pairs per CONDITION against the NULL. Error bars represent 95% confidence intervals.

### Results and discussion

We conducted a 4×3 mixed effect ANOVA with item TYPE (Rp+, Rp-, Nrp+, Nrp-) as a within-subject variable and CONDITION (S-S-S vs. S-T-T vs. S-S-T) as a between-subject variable on the final recall performance. Item TYPE had a significant main effect on recall performance, *F*(3,262) = 11.80, *p* < .001, *MSE* = 176.71, η^2^_p_ = .12, additionally, we also observed the main effect of CONDITION, *F*(2,87) = 4.17, *p* = .02, *MSE* = 546.26, η^2^_p_ = 0.09. The interaction between TYPE and CONDITION did not reach significance, *F*(6,261) = 1.48, *p* = .18, *MSE* = 176.71. BANOVA confirmed the main two effects model with a Bayes factor of 9.4×10^4^ in comparison to the null model. Bayes factors for other models against null can be seen on table 5. However, we only have a very week evidence against the model containing only the effect of item TYPE in favour of the previously mentioned model, *BF* = 2.37. The posterior likelihood of the two main effects model is decisively greater than the other possible models’ (e.g. the ones containing interaction), therefore we can exclude the emergence of interaction, see Table 6.

**Table 5.** Bayes factors for each possible BANOVA model for Experiment 1 AGAINST NULL model.

| Model | Bayes factor |
| --- | --- |
| **Type + Condition + Participant** | **$9.44 \times 10^4$** |
| **Type + Participant** | **$3.97 \times 10^4$** |
| Type + Condition + Type×Condition + Participant | $1.33 \times 10^4$ |
| Type + Type×Condition + Participant | $5.47 \times 10^3$ |
| Condition + Participant | 2.26 |
| Condition + Type×Condition + Participant | 0.23 |
| Type×Condition + Participant | 0.99 |

**Table 6.**
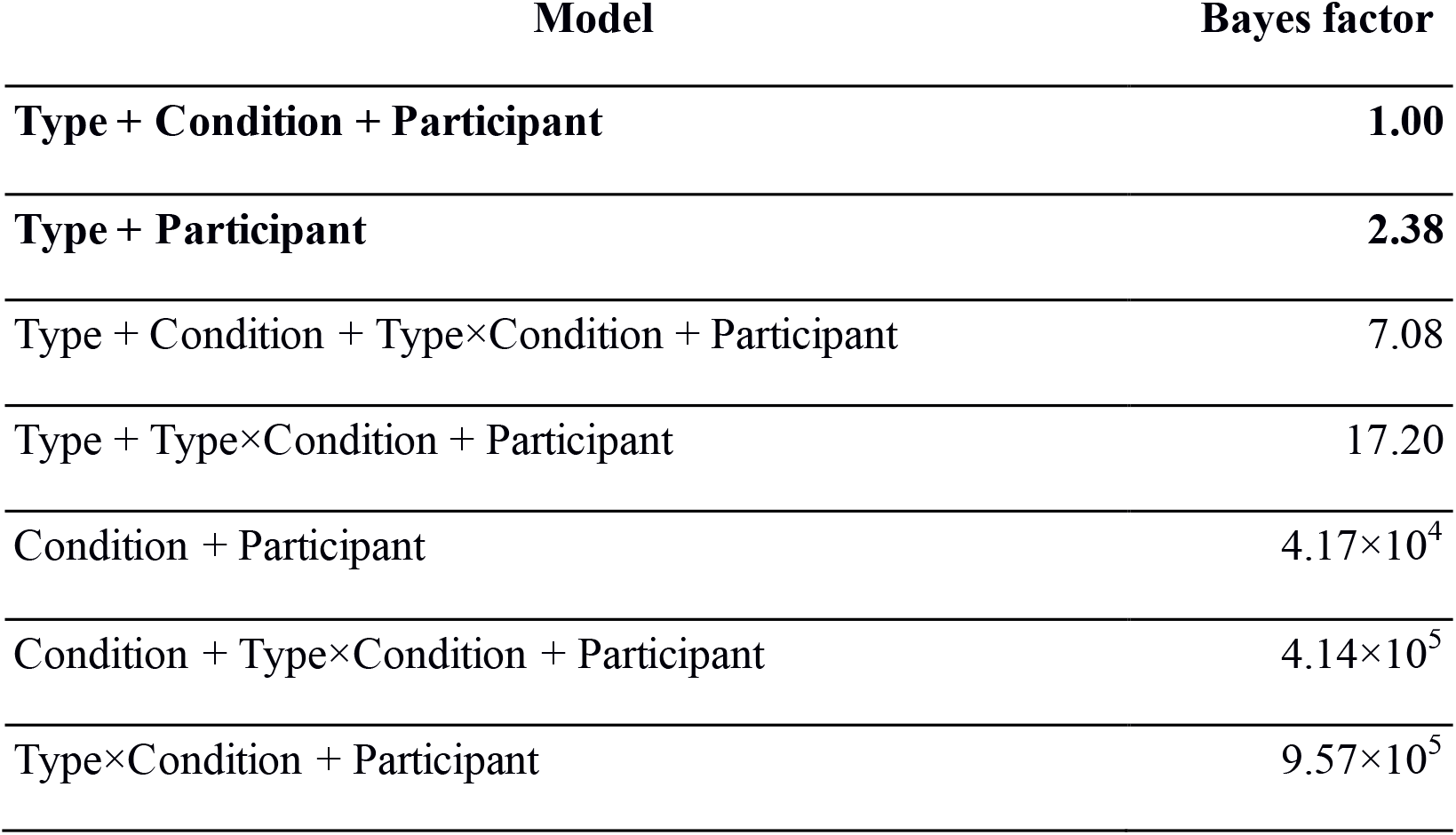
Bayes factors for each possible BANOVA model for Experiment 2 IN FAVOUR OF THE STRONGEST model.

To break down the main effect of item TYPE we conducted contrasted GLHT with Holm correction and paired Bayesian t-tests, as in Experiment 1. We did not observe any discrepancy between Rp- and Nrp- items: *t*(261) = −1.358, p = .494, *β* = −1.35 [−5.81, 3.10]. Bayesian t-test confirmed that there is no difference between the recall rates with a Bayes factor of 0.143 (which means *BF* = 6.99 in favour of the null hypothesis). Only the condition-independent practice effect was significant: *t*(261) = 11.235, *p* < .001, *β* = 11.23 [6.78, 15.69] with a Bayes factor of 3.5×10^5^. To investigate the main effect of CONDITION we conducted pairwise GLHT with Holm correction and independent samples Bayesian t-tests. The difference between S-S-S and S-S-T conditions approaches significance, *t*(87) = −6.76, *p* = .055, *β* = −6.76 [−13.95, 0.43], however *BF* = 12.2 endorses the inequality between group performances. We did not observe difference between performance on S-S-S and S-T-T conditions, *t*(87) = 1.39, *p* = .646, *β* = 1.39 [−5.80, 8.58] with a Bayes factor of 0.167 (BF = 5.99 in favour of the equality of recall rates). The difference between S-S-T and S-T-T groups was strongly confirmed, *t*(87) = 8.15, *p* = .025, *β* = 8.14 [0.95, 15.34] with a Bayes factor of 293.

To confirm the absence of RIF and practice effects in each conditions we conducted Bayesian t-tests within all groups separately. The analysis confirmed that retrieval-induced forgetting did not occur in any of the conditions, however the selective retrieval practice did not have a confirmable effect on Rp+ items within the S-S-T condition (see Table 7).

**Table 7.** Bayes factors for practice and RIF effects within each group in Experiment 2 against the absence of the effect.

| Condition | Bayes factor |  |
| --- | --- | --- |
|  | Practice | RIF |
| **S-T-T** | 12.64 | 0.23 (4.35 in favour of null) |
| **S-S-S** | 922.33 | 0.246 (4.07 in favour of null) |
| **S-S-T** | 1.00 | 0.207 (4.83 in favour of null) |

With the increased memory strength – as a consequence of any combination of full restudy and retest cycles – we can confirm the presence of a significant practice effect (except in restudy - full test condition on subgroup level, however we did not observe any kind of interaction between item TYPE and CONDITION) while the adverse memory effect of practice cannot be observed neither on model nor on subgroup level. Seemingly an extra study followed by a full cued recall test is an extremely efficient form of learning resulting high level of retention: the strength of representations shields the baseline items against the interference suppressing the practice effect, in comparison to other conditions.

### General discussion

In two experiments we manipulated the initial rate of learning in a retrieval-induced forgetting paradigm. Our results suggest that including a cued recall test for the whole study material following the initial study phase reduces retrieval-induced forgetting and leads to relatively lower target strengthening. The observed reduction in the practice effect by the initial test group was probably caused by the increased recall rate on baseline items. To further investigate the reduction of retrieval-induced forgetting, we performed regression analysis with overall recall performance and the amount of practice as predictors and RIF scores (Nrp- - Rp- differences) as outcome. We found a negative correlation between average rate of learning and differential RIF scores. Furthermore, the positive predictive strength of the observed practice effect was modulated by overall recall performance independently from the experimental manipulation. Additionally, after an increased level of global representational strength were induced with either repeated retrieval attempts or reexposures of stimuli we confirmed the total absence of retrieval-induced forgetting. More importantly, this increase in strength was independent from the magnitude of the practice effect. In the followings, we assume that the practice effect – the difference between performance on practiced items and their baseline – is a consequence of repeated selective retrieval, while representational strength – mentioned also as the rate of learning – refers to the learning success of the whole learning episode.

Based on the results of our first experiment, we extended the explanation of Racsmány & Keresztes (2015). First, we observed a crucial consequence of retrieval-based learning which was not handled in the previous models: initial retrieval shields the baseline items tested in the second block (Nrp+) against output interference resulting a decrement in the observed practice effect (for a detailed explanation see Szpunar, McDermott, & Roediger III, 2008). Results from Racsmány & Keresztes (2015) suggest that inserting a full initial cued recall before the selective retrieval practice leads to the lack of retrieval-induced forgetting. However, their practice method – in contrast with our experimental procedure – contained reexposure blocks after every cycles - might result a possible decrement in the level of competition caused by the selective retrieval during the second and third retrieval practice cycles.

Several previous studies investigated the influence of competitor and target strengthening separately (e.g. Anderson et al., 1994; Campbell & Phenix, 2009; Norman et al., 2007). However, none of these studies explored the effect after the entire set of studied items was simultaneously weakened or strengthened. Together with the results of Experiment 2 we extend the predictions of the neural network model of retrieval-induced forgetting on competitor- and target strengthening (see Norman et al., 2007). Our findings on the negative effect of learning rate arises many questions and possibly a demand for a revision about the key features of retrieval-induced forgetting. It has been demonstrated that retrieval-induced forgetting is not completely independent from target strength, however, their relation is not monotonic (Campbell & Phenix, 2009; Keresztes & Racsmány, 2013; Norman et al., 2007). Our results suggest that rate of learning and retrieval-induced forgetting have a direct negative relationship (see the results of Experiment 1). Additionally, the target strength-dependent nature of RIF can only be observed at a specific, middle-low-range level of overall encoding success. We also demonstrated that RIF can be eliminated by increasing representational strength. We can integrate our finding into the existing frameworks concluding that the rate of learning and retrieval-induced forgetting are related on a non-monotonic way. It can be forecasted – and derived from the predicted competitor weakening –, that RIF can not be expected on a low level of representational strength and here has been proved the total absence of the phenomenon on a high rate of learning.

These findings suggest that boundary conditions and key features of retrieval practice paradigm show varying characteristics at different representational strength of previously studied items. Consequently, retrieval-induced forgetting is rather one possible scenario following selective retrieval practice, which can be observed only at a moderate rate of learning, than a universal phenomenon.

## Acknowledgements

We thank Péter Pajkossy for his help in data analysis and Bertalan Polner for his comments on a previous version of the manuscript.

## Footnotes

1 Reversed practice is identical to practice method described in Anderson et al (1994), except that participants has to fill in the missing letters in the category cue instead of the exemplar (e.g. Fr___– Orange).

## Notes

* **Author note**: This work was supported by the National Brain Research Program of the Hungarian Academy of Sciences (KTIA_NAP_13-2-2014-0020 and 2017-1.2.1-NKP-2017-00002) and the NKFI K124094 Research Grants.

